# Structure elucidation of colibactin

**DOI:** 10.1101/574053

**Authors:** Mengzhao Xue, Chung Sub Kim, Alan R. Healy, Kevin M. Wernke, Zhixun Wang, Madeline C. Frischling, Emilee E. Shine, Weiwei Wang, Seth B. Herzon, Jason M. Crawford

**Affiliations:** Department of Chemistry, Yale University, New Haven, Connecticut 06520, United States; Chemical Biology Institute, Yale University, West Haven, Connecticut 06516, United States; Department of Microbial Pathogenesis, Yale School of Medicine, New Haven, Connecticut 06536, United States; Department of Molecular Biophysics and Biochemistry, Yale University School of Medicine, P.O. Box 208114, New Haven, Connecticut 06520, United States; W. M. Keck Biotechnology Resource Laboratory, Yale University School of Medicine, 300 George Street, New Haven, Connecticut 06510, United States; Department of Pharmacology, Yale School of Medicine, New Haven, Connecticut 06520, United States

## Abstract

Colibactin is a gut microbiome metabolite of unknown structure that has been implicated in colorectal cancer formation. Several studies now suggest that the tumorgenicity of colibactin derives from interstrand cross-linking of host DNA. Here we use a combination of genetics, isotope labeling, tandem MS, and chemical synthesis to deduce the structure of colibactin. Our structural assignment accounts for all known biosynthetic data and suggests roles for the final unaccounted enzymes in the colibactin gene cluster. DNA cross-link degradation products derived from synthetic and natural colibactin were indistinguishable by tandem MS analysis, thereby confirming the structure prediction. This work reveals the structure of colibactin, which has remained incompletely defined for over a decade.

## Introduction

Colibactin is the product of the *clb* (also referred to as *pks*) biosynthetic gene cluster (BGC) that is commonly present within *E. coli* found in the human colon (*1, 2*). Intensive interest in colibactin has been fueled by reports that *clb*^+^ *E. coli* induce DNA damage in eukaryotic cells in vitro (*3*) and in vivo (*4*), promote tumor formation in mouse models of colorectal cancer (CRC) (*5–7*), and are more prevalent in CRC patients than healthy subjects (*5, 8*). However, despite intensive efforts, the structure of colibactin has remained unknown for over a decade (*9–13*).

Traditionally, microbial natural products have been produced by large-scale fermentation of bacterial cultures, discovered via activity-guided isolation, and their structures determined by well-established chemical spectroscopic methods. However, because colibactin has been recalcitrant to these classical isolation techniques, our knowledge of its structure and biological activity is derived from diverse interdisciplinary findings. Evaluating the biological effects of *clb*^+^ bacterial strains has revealed colibactin induces DNA damage in eukaryotic cells. Enzymology, bioinformatic analysis of the *clb* BGC, stable isotope feeding experiments, characterization of biosynthetic intermediates, and gene deletion and editing studies have given insights into many elements of colibactin’s biosynthesis, biological activity, and cellular trafficking. Additionally, by employing chemical synthesis to access shunt metabolites and putative biosynthetic intermediates, a mechanism of action model has been developed that defines the key structural elements of colibactin underling its DNA-damaging properties.

Merging of this data forms a picture, albeit incomplete, of colibactin’s biosynthesis, structure, and mode of genotoxicity. Colibactin is assembled in a linear prodrug form referred to as precolibactin (see **1**, Fig. 1). Key structural elements of precolibactins include a terminal *N*-myristoyl-d-Asn amide (blue in **1**) (*14–16*) and an aminocyclopropane residue (green in **1**) (*17–19*). The terminal amide is cleaved in the periplasm by the pathway-dedicated serine protease, colibactin peptidase (ClbP) (*20, 21*). The resulting amine **2** undergoes a series of spontaneous cyclization reactions to generate spirocyclopropyldihydro-2-pyrrolones resembling **3** (*22, 23*). These cyclizations place the cyclopropane in conjugation with both an imine and amide, rending the cyclopropane electrophilic and now capable of alkylating DNA (*12, 22*). A recent study established that *clb*^+^ *E. coli* cross-link DNA (*24*), suggesting that colibactin contains a second electrophilic site that has yet to be elucidated. While the colibactin–adenine adduct **4** has been characterized independently by two different laboratories (*25, 26*), the full structure of colibactin and the site of the second alkylation have remained undefined.

**Fig. 1.**
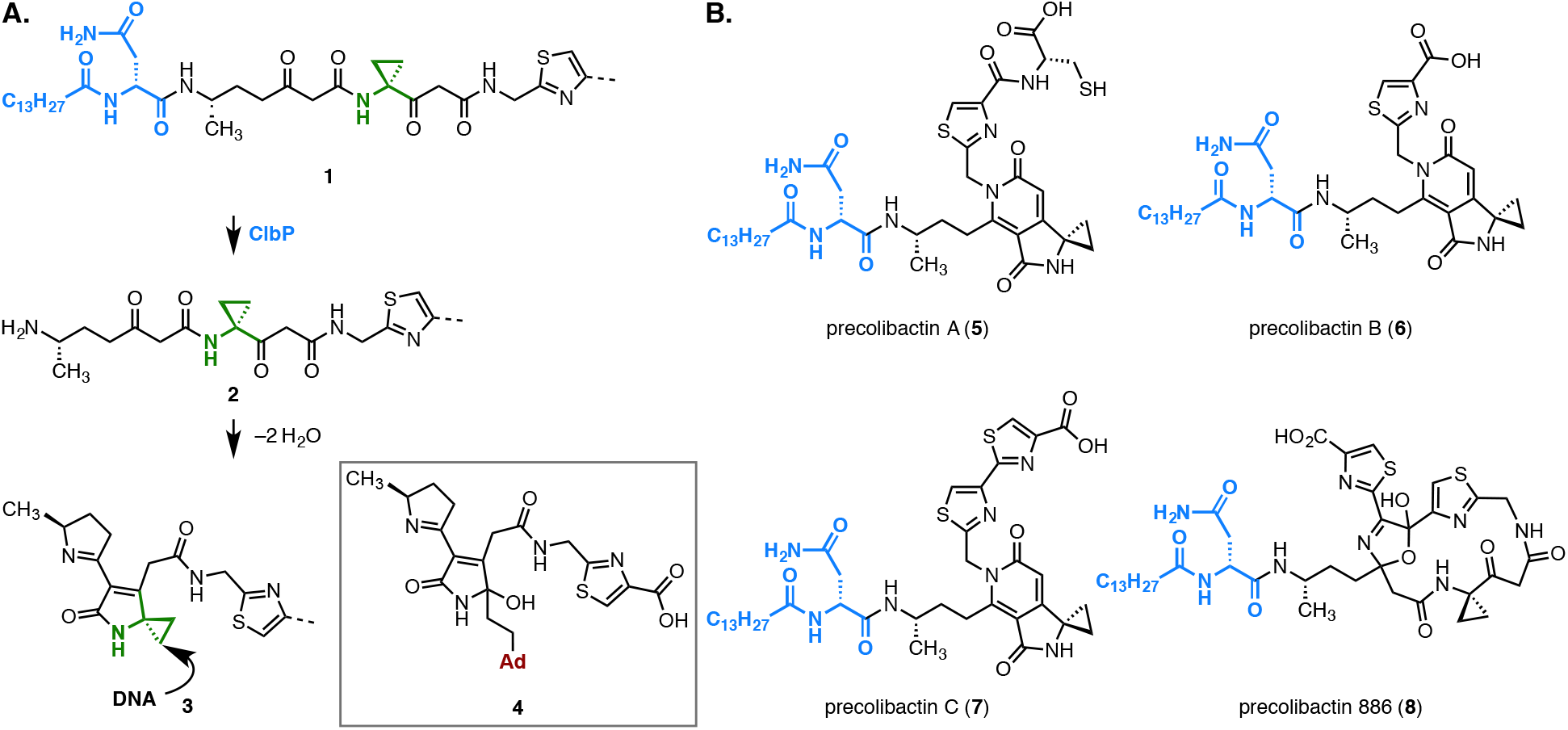
**A.** Mechanism of DNA alkylation by *clb* metabolites formed in wild-type cultures. **B.** Structures of *clb* metabolites formed in mutant Δ*clbP clb*^+^ *E. coli* cultures.

Precolibactins A–C (**5–7**) and precolibactin 886 (**8**), were identified in cultures of unnatural *clbP* mutants (*16, 17, 19, 27–30*). This genetic modification was introduced to facilitate isolation efforts. Unwittingly, it also changes the fate of the linear precursor **1**, resulting in pyridone formation (for **5–7**) (*12, 22*) or macrocyclization (for **8**) (*31*). Precolibactin 886 (**8**) requires every biosynthetic gene in the pathway except polyketide synthase (PKS) *clbO*, type II thioesterase *clbQ*, and peptidase *clbL* (*23*). Early genetic studies established that deletion of any biosynthetic gene in *clb* abolishes cytopathic effects (*3*), thus (pre)colibactin is believed to possess additional chemical functionalities not represented in **8**. Recently a derivative of precolibactin 886 (**8**) bearing a terminal oxazole ring was reported (*32*).

### Characterization of colibactin–DNA crosslinks and biosynthetic proposal

Because colibactin has proven recalcitrant to isolation, we turned our efforts towards structural elucidation of the DNA cross-links generated by *clb*^+^ *E. coli* (*24*). By employing tandem MS analysis of the digestion mixture containing linearized pUC19 DNA that had been exposed to *clb*^+^ *E. coli* (*25*) the colibactin–adenine adduct **4** was recently identified. In that study, wild-type *E. coli* BW25113 and its cysteine and methionine auxotrophs (Δ*cysE*, Δ*metA*) (*33*) containing *clb* on a bacterial artificial chromosome (BAC) were employed. The latter two cultures were supplemented with L-[U-^13^C]-Cys or L-[U-^13^C]-Met, which are known precursors to the thiazole (*17*) and aminocyclopropane (*17, 18, 34*) residues of colibactin, respectively. This approach allowed for the identification of *clb* metabolite–nucleobase adducts by mining for isotope shifts in the mass spectra from unlabeled wild-type and labeled auxotrophic cultures.

Further analysis of this data revealed a compound of *m/z* = 537.1721 (*z* = 2, Fig. 2A). This mass corresponds to a molecular formula of C_47_H_50_N_18_O_9_S_2_^2+^ (error = 0.37 ppm). The doubly-charged ion was shifted by +3 or +4 units in cultures containing L-[U-^13^C]-Cys or L-[U-^13^C]-Met, respectively (cysteine auxotroph: C_41_^13^C_6_H_50_N_18_O_9_S_2_^2+^, *m/z* 540.1821, error = 0.28 ppm; methionine auxotroph: C_39_^13^C_8_H_50_N_18_O_9_S_2_^2+^, *m/z* 541.1857, error = 0.74 ppm; Figs. 2B, 2C, respectively), supporting the presence of two thiazole and two cyclopropane residues.

**Fig. 2.**
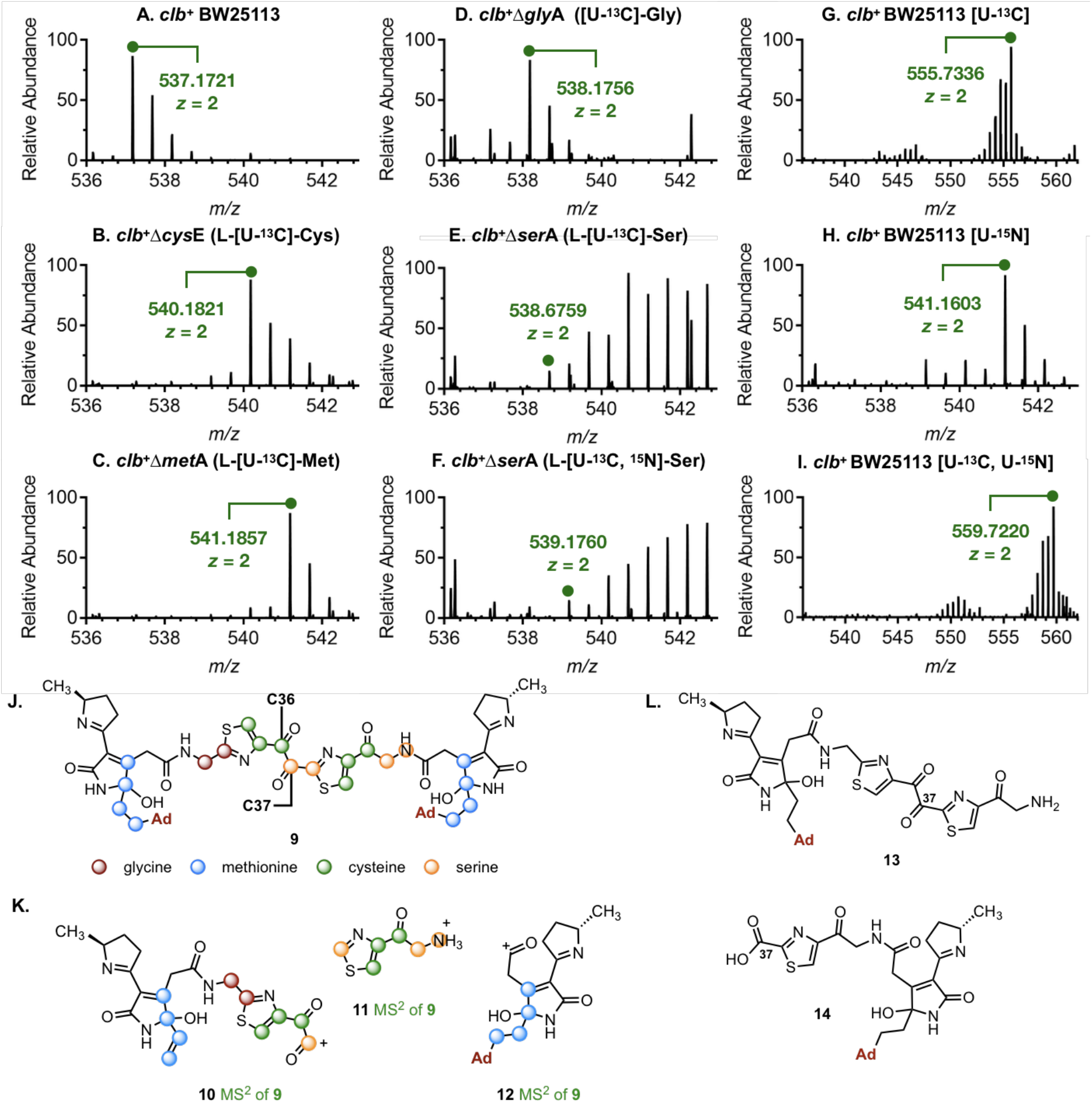
**A–I:** Selected HRMS signals deriving from treatment of linearized pUC19 DNA with *clb*^+^ *E. coli*, followed by digestion. **A. 9** extracted from *clb*^+^ BW25113and M9-casamino acids (CA) medium (*z* = 2, theoretical = 537.1719, experimental = 537.1721, error = 0.37 ppm). **B.** [^13^C_6_]-**9** extracted from *clb*^+^Δ*cysE* and M9-CA medium containing L-[U-^13^C]-Cys (*z* = 2, theoretical = 540.1820, experimental = 540.1821, error = 0.28 ppm). **C.** [^13^ C8]-9 extracted from *clb*^+^Δ*metA* and M9-CA medium containing L-[U-^13^C]-Met (*z* = 2, theoretical = 541.1853, experimental = 541.1857, error = 0.74 ppm). **D.** [^13^C_2_]-**9** extracted from *clb*^+^Δ*glyA* and M9-CA medium containing [U-^13^C]-Gly (*z* = 2, theoretical = 538.1753, experimental = 538.1756, error = 0.65 ppm). **E.** [^13^C_3_]-**9** extracted from *clb*^+^Δ*serA* and M9-CA medium containing L-[U-^13^C]-Ser (*z* = 2, theoretical = 538.6769, experimental = 538.6759, error = 1.90 ppm. **F.** [^13^C_3_, ^15^N]-**9** extracted from *clb*^+^Δ*serA* and M9-CA medium containing L-[U-^13^C, ^15^N]-Ser (*z* = 2, theoretical = 539.1754, experimental = 539.1760, error = 1.07 ppm). **G.** [^13^C_37_]-**9** extracted from *clb*^+^ BW25113 and M9-glucose medium containing D-[U-^13^C]-glucose (*z* = 2, theoretical = 555.7339, experimental = 5555.7336, error = 0.49 ppm). **H.** [^15^N_8_]-**9** extracted from *clb*^+^ BW25113 and M9-glucose medium containing ^15^NH_4_Cl (*z* = 2, theoretical = 541.1599, experimental = 541.1603, error = 0.74 ppm). **I.** [^13^C_37_, ^15^N_8_]-**9** extracted from *clb*^+^ BW25113 and M9-glucose medium containing D-[U-^13^C]-glucose and ^15^NH_4_Cl (*z* = 2, theoretical = 559.7219, experimental = 559.7220, error = 0.22 ppm). **J.** Structure of the colibactin–bis(adenine) adduct **9. K.** Structures **10–12** are daughter ions and tandem MS fragments supporting the structure of **9. L.** The known adenine adduct **4** (see Fig. 1A; *z* = 1, theoretical = 540.1772, experimental = 540.1777, error = 0.93 ppm) and the two novel DNA adducts **13** (*z* = 2, theoretical = 346.5945, experimental = 346.5944, error = 0.29 ppm) and **14** (*z* = 1, theoretical = 568.1721, experimental = 568.1716, error = 0.88 ppm) were also detected in the DNA digestion mixtures. Detection of **13** explains why *clbL* mutants alkylated exogenous DNA but could not crosslink DNA.

To gain further insights into the structure, we analyzed its production in glycine (Δ*glyA*) and serine (Δ*serA*) auxotrophs. These cultures were supplemented with [U-^13^C]-Gly (for the glycine auxotroph), L-[U-^13^C]-Ser, or L-[U-^13^C, ^15^N]-Ser (for the serine auxotroph). Glycine serves as the CN extension in the 2-methylamino thiazole of precolibactin A (**5**) (*17*), whereas serine is incorporated into precolibactin-886 (**8**) via an unusual α-aminomalonate extender unit (*29, 30, 35*). The doubly-charged ion was shifted by 1 unit in cultures containing [U-^13^C]-Gly (C_45_^13^C_2_H_50_N_18_O_9_S_2_^2+^, *m/z* 538.1756, error = 0.65 ppm; Fig. 2D), suggesting incorporation of one glycine building block, as expected. However, the doubly-charged ion was shifted by 1.5 units in cultures containing L-[U-^13^C]-Ser (C_44_^13^C_3_H_50_N_18_O_9_S_2_^2+^, *m/z* 538.6759, error = 1.90 ppm), and by two units (1.5 units from carbon mass difference and 0.5 unit nitrogen mass difference) in cultures containing L-[U-^13^C, ^15^N]-Ser (C_44_^13^C_3_H_50_N_17_^15^NO_9_S_2_^2+^, *m/z* 539.1760, error = 1.07) unexpectedly indicating three carbon atoms and one nitrogen atom are derived from serine (Figs. 2E, 2F, respectively). We note that these cultures produced a range of higher molecular weight isotopologs owing to amino acid metabolism and incorporation into other building blocks (see below).

Additional cultures of wild-type *clb*^+^ *E. coli* grown in minimal medium lacking amino acids and supplemented with D-[U-^13^C]-glucose were analyzed. In these cultures, the doubly-charged ion was shifted by 18.5 units (C_10_^13^C_37_H_50_N_18_O_9_S_2_^2+^, *m/z* 555.7336, error = 0.49 ppm; Fig. 2G), establishing the colibactin residue contained 37 carbon atoms. Cultivation in minimum medium containing [^15^N]-ammonium chloride shifted the doubly-charged ion by 4 units (C_47_H_50_N_10_^15^N_8_O_9_S_2_^2+^, *m/z* 541.1602, error = 0.55 ppm; Fig. 2H), indicating the colibactin residue contained eight nitrogen atoms. A double-labeling experiment using D-[U-^13^C]-glucose and [^15^N] ammonium chloride resulted in a shift of the doubly-charged ion by 22.5 units (17.5 units from carbon mass difference, and 4 units from nitrogen mass difference), confirming the results of the individual labeling experiments (C_10_^13^C_37_H_50_N_10_^15^N_8_O_9_S_2_^2+^, *m/z* 559.7220, error = 0.22 ppm; Fig. 2I). The mass of the non-labeled doubly-charged ion and the results of the labeling experiments suggested a molecular formula of C_37_H_40_N_8_O_9_S_2_ for the colibactin fragment. An ion corresponding to protonated adenine was detected in the tandem MS, while the neutral mass loss of two consecutive adenine bases and two hydroxyl groups were observed in all labeling experiments. Finally, the singly-charged (*z* = 1) and triply-charged (*z* = 3) ions corresponding to this structure was also detected in many of these auxotrophs and provided data of comparable quality (Figs. S45, S49–S51 and Tables S1–S9).

Based on these data, we reconsidered the possible unaccounted functions of ClbO, ClbQ, and ClbL (Fig. 3). ClbO is a polyketide synthase that accepts an a-aminomalonyl-extender unit in protein biochemical studies (*30*), suggesting a canonical polyketide-type extension step. ClbQ serves as an editing thioesterase and also off-loads intermediary structures (*23, 29, 36*). While it is well-known that these off-loaded structures enhance the metabolite diversity arising from the pathway, we reasoned that they could also serve as downstream substrates. In the absence of a dedicated terminal thioesterase, we considered ClbL’s role in interacting with ClbO and ClbQ products in the final enzymatic steps of precolibactin biosynthesis (prior to ClbP-mediated deacylation). We reasoned that an uncharacterized ClbL-mediated transpeptidase activity could promote a heterodimerization event employing a structure that had been off-loaded by ClbQ. Such a structure would accommodate the isotopic labeling studies, including the presence of two aminocyclopropane units derived from methionine (Fig. 3, and see below).

**Fig. 3.**
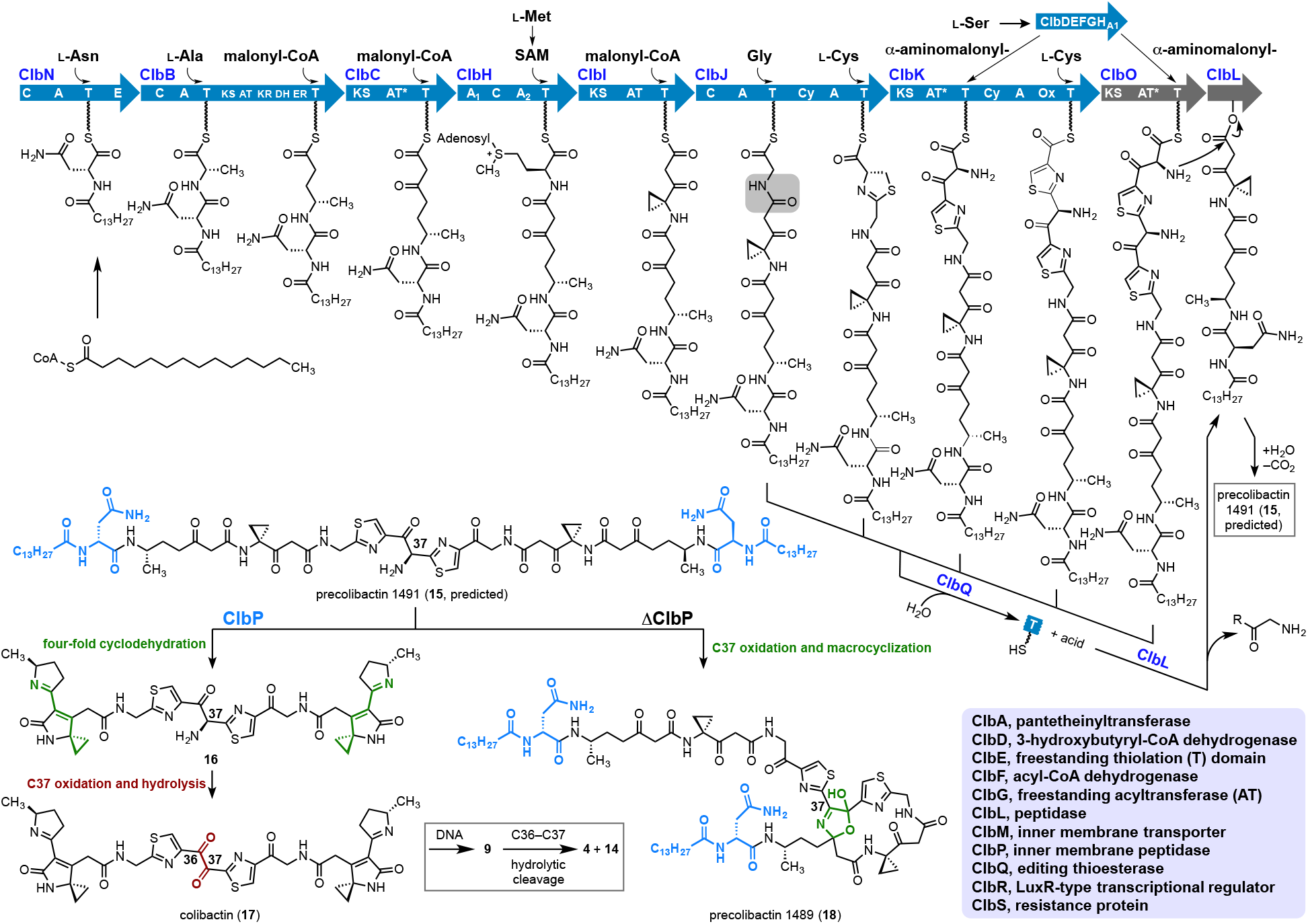
Proposed biosynthesis of (pre)colibactin. Precolibactin is assembled through a modular hybrid nonribosomal peptide synthetase (NRPS)–polyketide synthase (PKS) pathway. Based on our isotopic labeling studies, we predict the functions of the unaccounted PKS ClbO and transpeptidase ClbL (grey), and we provide a new rationale for why editing thioesterase ClbQ offloads truncated products and is required for genotoxicity. In this proposal, ClbO extends the chain through a canonical PKS event using an a-aminomalonyl-extender unit. ClbQ serves an editing function and also offloads intermediary structures by hydrolyzing acyl- and peptidyl-carrier protein thioesters [represented as thiolation (T) domains]. Hydrolysis products incorporating glycine could serve as ClbL transpeptidase substrates to regulate precolibactin heterodimerization. Precolibactin is activated by peptidase ClbP, to liberate two *N*-myristoyl-d-Asn prodrug residues and colibactin, which undergoes spontaneous cyclodehydration sequences to generate two warheads. In the absence of ClbP, precolibactin 1489 (**18**) forms through an alternative macrocyclization sequence. Amino acids are depicted at their sites of pathway entry and are supported by isotopic labeling studies. Additional domain abbreviations: C, condensation; A, adenylation; E, epimerization; KS, ketosynthase; KR, ketoreductase; DH, dehydratase; ER, enoylreductase; AT*, inactivated acyltransferase (AT); Cy, dual condensation/cyclization; Ox, oxidase.

Considering the isotope labeling and MS data in the context of these biosynthetic hypotheses, we formulated the structure of the observed parent ion as the bis(adenine) adduct **9** (Fig. 2J). The experimental and theoretical masses are in agreement (537.1719 and 537.1721, respectively; *z* = 2; error = 0.37 ppm). The positions of the cysteine, methionine, serine, and glycine isotopic labels depicted in **9** are fully supported by all known elements of colibactin biosynthesis and tandem MS analysis (Figs. S1–S3, S74–S80 and Tables S10, S13, S17, S21, S25, S29, S34, S38, S42). The tandem MS fragments **10–12** shown in Fig. 2K provide further robust support for the structure **9**. Each of the ions **10–12** possessed the expected mass shift in the individual labeling experiments (Figs. S74–S80).

The structure **9** is also in agreement with the expected chemical reactivity of colibactin. Functionally, the bis(adenine) adduct **9** derives from two-fold alkylation of DNA, which is consistent with the earlier determination that simpler colibactins alkylate DNA by nucleotide addition to the cyclopropane (*22*), the isolation of the colibactin–adenine adduct **4** (*25, 26*) and, importantly, the observation that *clb*^+^ *E. coli* cross-link DNA and activate cross-link repair machinery in human cells (*24*). In agreement with the known propensity of α-diketones to hydrate under aqueous conditions (the *K*_d_ for dissociation of the monohydrate of butane-2,3-dione = 0.30) (*37*), the product of hydration of C37 (**S1**) was also detected (Figs. S45–S51 and Tables S1–S9). Tandem MS and isotopic labeling data for **S1** fully support the structure of the hydrate and are in agreement with the diketone form **9** (Figs. S81–S88 and Tables S10, S14, S18, S22, S26, S30, S35, S39, S43). The aminal functional groups in **9** derive from aerobic oxidation of the ring-opened products, as previously established (*25, 38*).

Additional nucleobase adducts were detected at discrete retention times (Fig. 2L). We detected the known adduct **4** (Fig. 1A)(*25, 26*), the methylaminoketone **13**, and the right-hand fragment **14** in the digestion mixtures. We have demonstrated that that the C36–C37 bond in advanced colibactins (see **9** for numbering) is susceptible to nucleophilic cleavage (*31*). Hydrolytic degradation of **9** at this bond accounts for isolation of the earlier monoadenine adduct **4** (*25, 26*) and now, the right-hand fragment **14**. The detection of **13** explains why *clbL* mutants could alkylate but not crosslink exogenous DNA in prior studies (see also Fig. 3) (*39*).

### Identification of colibactin (17)

Based on these data, we searched *clb*^+^ *E. coli* cultures for the structures of the natural colibactin **16**, the corresponding a-ketoimine oxidation product (not shown), and the α-dicarbonyl **17** (Fig. 3). While **16** and its oxidation products (half-life unknown) were not detected under the conditions of our experiments, we detected both the proton and sodium adducts of **17** in *E. coli* DH10B harboring the *clb* BAC ([M+H]^+^ theoretical = 771.2378, experimental 771.2367, error = 1.47 ppm; [M+Na]^+^ theoretical = 793.2197, experimental = 793.2176, error = 2.64 ppm). Importantly, the metabolite was abrogated in a *clbO* deletion mutant, a *clbL* active site point mutant (S179A), and a *clb*^−^ BAC control strain (Fig. 4A). Because colibactin (**17**) was detected at low abundance, we also confirmed production in wild-type *E. coli* Nissle 1917 (Fig. 4B). Deletion of the *clb* genomic island (*16*) in Nissle 1917 destroyed production, as expected. To improve titers for robust isotopic labeling, we constructed a *clbS* colibactin resistance (*38, 40*) mutant in Nissle 1917. While deletion of *clbS* leads to a fitness defect and activates a clb-dependent bacterial SOS DNA damage response (*40*), this genetic modification resulted in an 8.5-fold improvement in the mass spectral signal intensity of colibactin (**17**). This is likely due to decreased processing of colibactin (**17**) by ClbS-mediated hydrolytic opening of the cyclopropane (*38*).

**Fig. 4.**
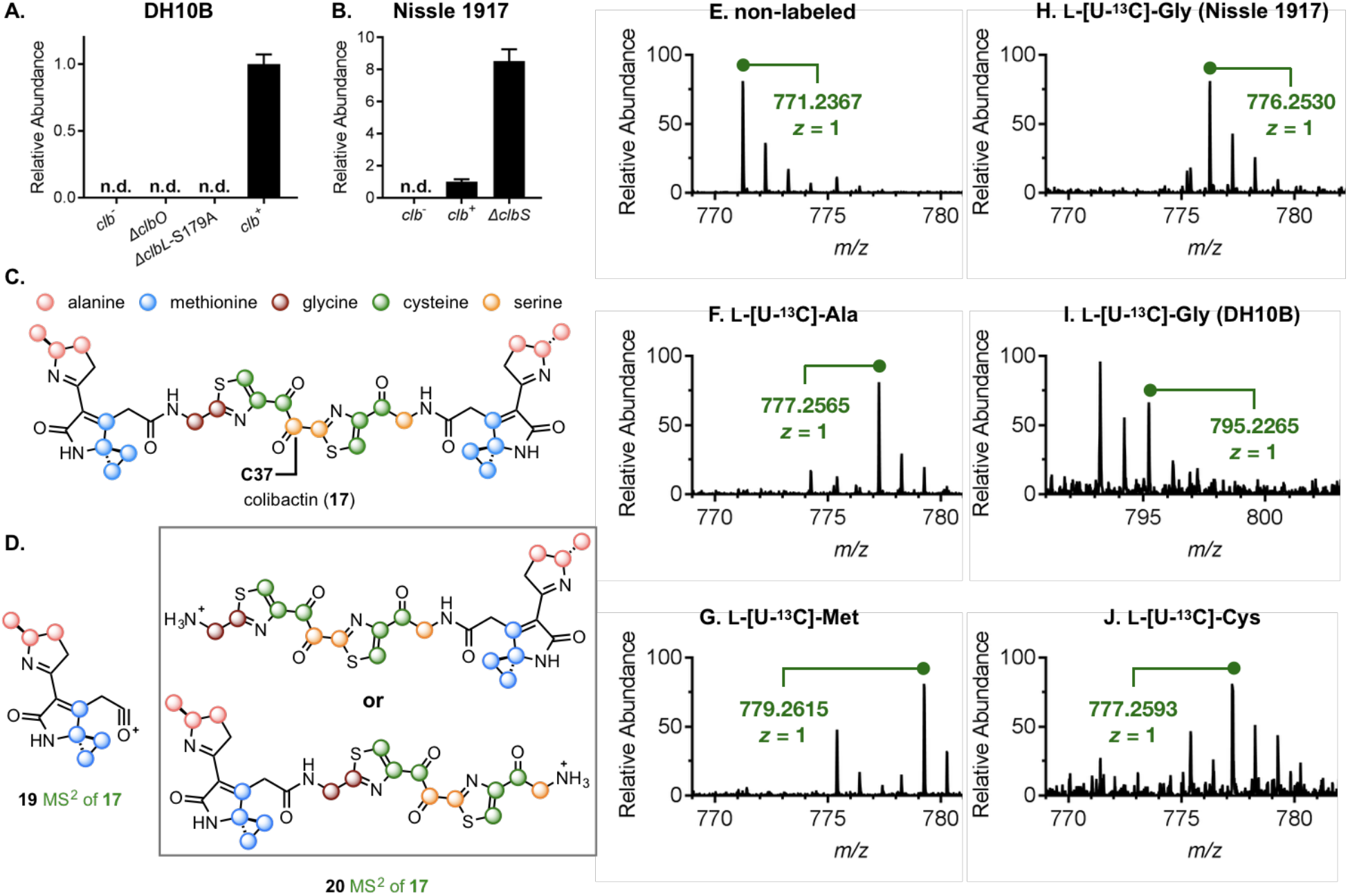
Stimulation, genetic dependence, and isotopic labeling of natural colibactin (17). **A.** Colibactin (**17**) was detected in *clb*^+^ DH10B cells, and deletion of the pathway (*clb*^−^), PKS *clbO*, and transpeptidase *clbL* (ClbL S179A) in a wild-type genetic background abrogated production. **B.** Colibactin (**17**) was detected in wild-type Nissle 1917, it was abrogated in a full *clb* genomic island deletion, and it was stimulated in a *clbS* colibactin resistance mutant. **C.** Proposed isotopic labeling pattern of colibactin (**17**). **D.** Major ions established from the tandem MS of **17**, including their proposed isotopic labeling patterns. The two structures of ion **20** are equally plausible based on the MS data. **E-J.** Results of isotopic labeling studies of **17** in Nissle 1917 (Δ*clbS*). [U-^13^C]-Gly labeling was conducted in both Nissle 1917 (Δ*clbS*)(**H**) and *clb*^+^ DH10B (**I**).

We proceeded with universal ^13^C-isotopic amino acid labeling studies in the bacterial strains. We individually supplemented *E. coli* Nissle 1917 (Δ*clbS*) cultures with labeled amino acids (200 mg/L). Colibactin (**17**) (Fig. 4C–J) incorporated two equivalents of Cys, Met, and Ala, as expected based on its proposed structure (Fig. 4E–G, J). The cultures labeled with L-[U-^13^C]-Ser and [U-^13^C]-Gly produced a range of isotopologs owing to their metabolism and incorporation into other building blocks (Fig. 4H, Fig. S10). To account for this variation, we used Ser-derived enterobactin, an iron-scavenging siderophore in *E. coli*, as an internal control for comparison (Fig. S10). We also repeated Gly labeling in *clb*^+^ DH10B for confirmation of dominant mono-labeling of Gly in colibactin (**17**) (Fig. 4I). We observed the tandem MS ions **19** and **20** of labeled and unlabeled colibactin (**17**), which are consistent with the proposed structure (Fig. 4D). We note that the two structures of **20** shown are equally plausible based on the MS data. Similar to the colibactin–DNA adducts, we observed the C37 hydrate of **17** (**S33**), which was supported by isotopic labeling and tandem MS (Figs. S12–S13).

### Characterization of precolibactin 1489 (18)

It was previously shown that every biosynthetic enzyme encoded in the *clb* gene cluster is required for cytopathic effects (*3*). While truncated precolibactins such as precolibactin 886 (**8**, Fig. 1B) can be detected as macrocyclization products in non-genotoxic *clbP* peptidase mutants (*29*), precolibactin 886 (**8**) is still produced in mutants of *clbL, clbO*, and *clbQ* in a *clbP*-deficient genetic background (ClbP S95A active site point mutant) (*23*). Additionally, a recently isolated precolibactin from a *clbP, clbQ*, and *clbS* triple mutant (*32*) does not account for *clbL* and *ClbQ* and was undetectable in freshly prepared organic extracts of a *clbP*-deficient strain under the conditions of our experiments. Given the structure of colibactin (**17**) and the requirement of every biosynthetic enzyme for cytopathic effects (3), we reasoned that more complex precolibactins existed.

To probe for this, we searched for the precolibactin that could account for colibactin (**17**) in the *clb*^+^ DH10B strain (ClbP S95A) (*23*). While we were not able to detect the expected unstable linear precursor precolibactin 1491 (**15**, Fig. 3) or its oxidation products under our extraction, processing, and analysis conditions, we were able to detect a metabolite that we predict to be the macrocycle precolibactin 1489 (**18**) (see Fig. 3, Fig. 5B, Fig. S15). We detected both the proton and sodium ion adducts of precolibactin 1489 (**18**; [M+H]^+^ theoretical = 1490.7786, experimental 1490.7773, error = 0.86 ppm; [M+Na]^+^ theoretical = 1512.7605, experimental = 1512.7565, error = 2.64 ppm). We evaluated the production of precolibactin 1489 (**18**) using our domain-targeted metabolomics approach (*23*). Briefly, we employed genome editing to individually inactivate the catalytic domains from ClbH to ClbL in the modular biosynthetic pathway (see Fig. 3). Importantly, precolibactin 1489 (**18**) was genetically dependent on all of the enzymatic steps of precolibactin biosynthesis (Fig. 5A). Production was only detected in an acyltransferase (AT) domain mutant of ClbI; metabolites dependent on this single domain can be complemented *in trans* by other ATs in the cell (*23*). Thus, precolibactin 1489 (**18**) represents the first reported product derived from the complete *clb* biosynthetic pathway.

**Fig. 5.**
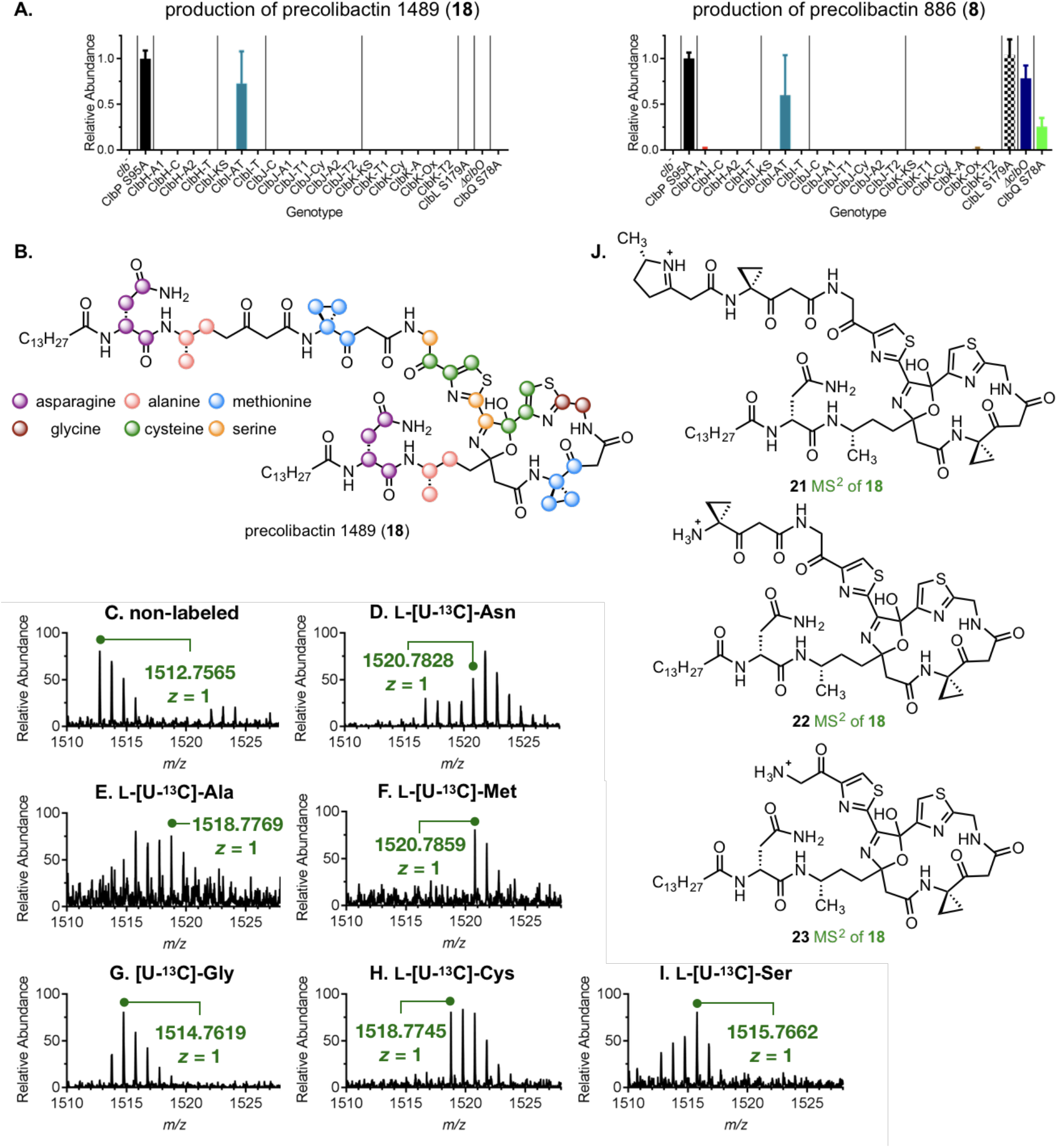
Genetic, tandem MS, and isotopic labeling support of proposed precolibactin 1489 (18). **A.** Individual inactivation of the catalytic domains between ClbH and ClbL in the biosynthetic pathway (see Fig. 3) through genome editing demonstrates that precolibactin 1489 (**18**) is dependent on all of the required final enzymatic steps of precolibactin biosynthesis (multidomain signatures are shown). Gene and domain designations are shown on the x-axis, and the abundance of precolibactin 1489 (**18**) in the specific mutants, relative to its production in Δ*clbP*, is shown on the *y*-axis. Precolibactin 886 (**8**) is not dependent on *clbL, clbO*, and *clbQ*, which are required for cellular genotoxicity (*3*). **B.** Proposed isotopic labeling pattern of precolibactin 1489 (**18**). **C–I.** Isotopic labeling studies of precolibactin 1489 (**18**) in a *clb*^+^ DH10B strain deficient in ClbP catalytic activity (ClbP-S95A). **J.** Proposed structures of ions **21–23** derived from the tandem MS of precolibactin 1489 (**18**).

To provide structural support for precolibactin 1489 (**18**), we conducted extensive ^13^C-isotopic amino acid labeling and tandem MS analysis (Fig. 5C–I and Fig. S14), as described above for colibactin (**17**, Fig. 4C) and the bis(adenine) adduct **9** (Fig. 2J). Labeled Met, Gly, Ala, Cys, and Ser precursors incorporated into precolibactin 1489 (**18**) in a manner consistent with its biosynthesis and structure. In addition to these colibactin substrates, two units of L-[U-^13^C]-Asn were incorporated, supporting the presence of two *N*-myristoyl-d-Asn residues.

Tandem MS analysis of precolibactin 1489 (**18**) produced the ions **21–23** and **S34–S36**, which are consistent with the proposed structure (Fig. 5J, Fig. S14C). Based on the recent determination that deacylation of precolibactin 886 (**8**) produces a non-genotoxic pyridone (*31*), it seems likely that precolibactin 1489 (**18**) is simply a stable product arising from oxidation and macrocyclization of the putative linear precursor precolibactin 1491 (**15**, Fig. 3). Regardless, these studies support a two-fold *N*-acyl-d-Asn prodrug activation mechanism, in which ClbP peptidase sequentially initiates the formation of two potent genotoxic architectures.

### Confirmation of the structure of colibactin (17)

The structure of colibactin (**17**) was confirmed through chemical synthesis. The presence of three reactive electrophilic sites (two spirocyclopropyldihydro-2-pyrrolones (*22*) and the hydrolytically-labile C36–C37 α-dicarbonyl (*31*)) necessitated a careful analysis of potential synthetic pathways. Fig. 6A outlines the essential elements of our strategy. While in earlier studies (*22*) monomeric colibactins were assembled by a linear approach (stepwise formation of bonds a, b, and c, in that order; see **24**, Fig. 6A) we recognized that colibactin (**17**) could be assembled by a two-fold coupling (a,a’ bond formation) of the diamine **26** with the novel β-ketothioester **25**. In addition to increased convergence, this approach masks the reactive (*22*) spirocyclopropyldihydro-2-pyrrolones as identical stable vinylogous imides. We have established that *N*-deacylation followed by mild neutralization is sufficient to induce cyclization and formation of the spirocyclopropyldihydro-2-pyrrolones residues (*22, 39*). Based on our previous observation that α-aminoketones at the C36–C37 position underwent spontaneous oxidation (*31*), we targeted an α-hydroxyketone in place of the α-dicarbonyl in colibactin (**17**). This was projected to allow for the assembly of **26** by benzoin addition, followed by late-stage oxidation to generate the α-dicarbonyl. Anecdotally, the initial ketone was generated at C36 in our synthesis of precolibactin 886 (**8**) (*31*). In exploratory experiments, we found that intermediates with a C37 ketone were more stable and pursued these, as outlined below.

**Fig. 6.**
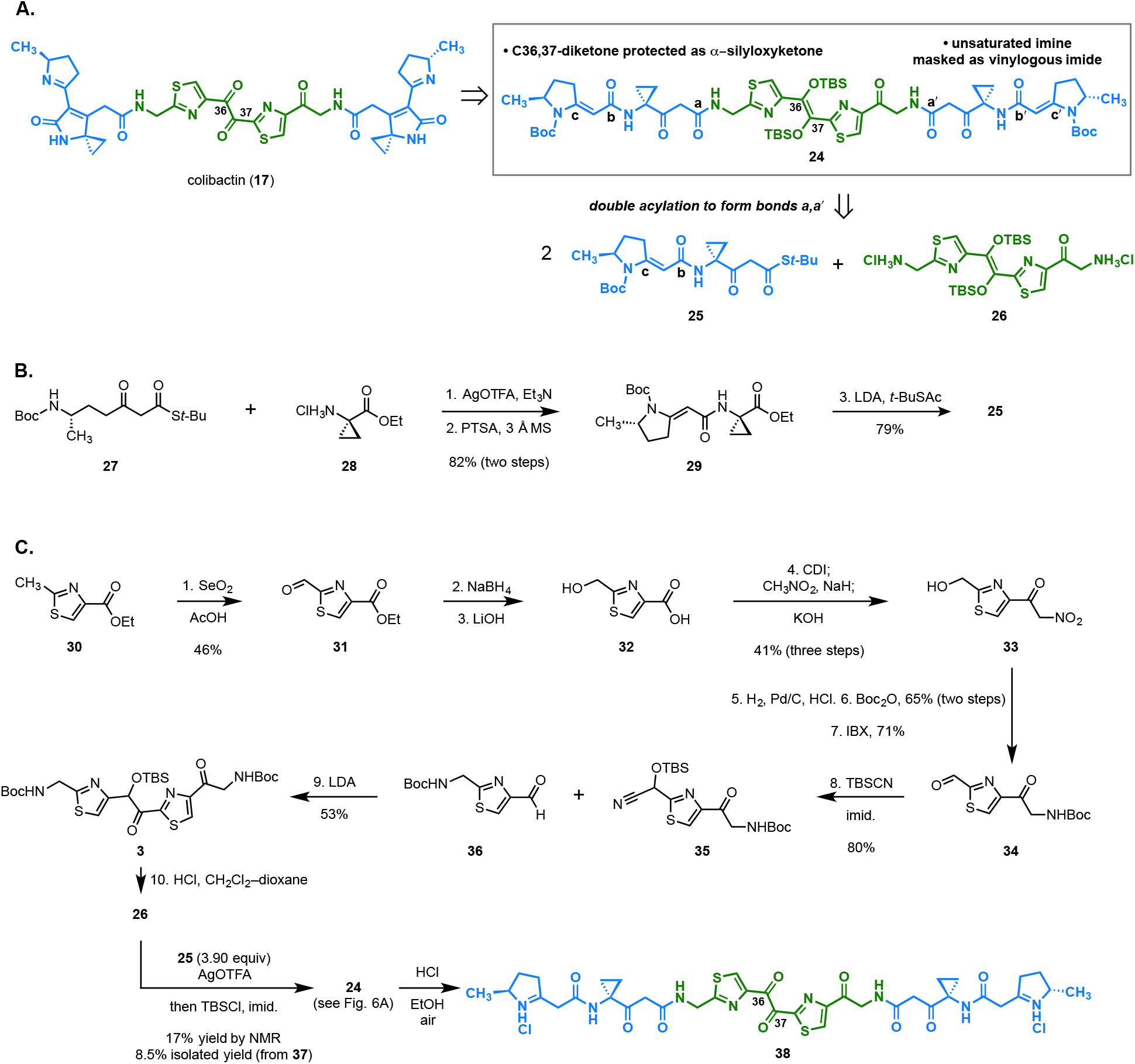
Synthesis of the colibactin precursor **38.**

The synthesis of the β-ketothioester **25** is shown in Fig. 6B. Silver trifluoroacetate-mediated coupling of the known β-ketothioester **27** (*22*) with ethyl 1-aminocyclopropyl-1-carboxylate **28** generated a linear coupling product (not shown) that was cyclized to the vinylogous imide **29** by treatment with *para*-toluenesulfonic acid (PTSA) in the presence of 3 Å molecular sieves (82%, two steps). Addition of the lithium enolate of *tert*-butyl thioacetate to **29** then generated the β-ketothioester **25** (79%).

The diamine **26** was synthesized by the route shown in Fig. 6C. Selenium dioxide oxidation of the commercial thiazole **30** generated the aldehyde **31** (46%). Reduction of the aldehyde (sodium borohydride), followed by saponification of the ester generated the hydroxy acid **32**. Treatment of the hydroxy acid **32** with excess 1,1′-carbonyldiimidazole (CDI) resulted in acylation of the primary alcohol and activation of the carboxylic acid as the expected acyl imidazole (LC/MS analysis). Addition of the sodium enolate of nitromethane, followed by in situ hydrolysis of the acylated alcohol, generated the β-nitroketone **33** (41% from **31**). Hydrogenolysis of the nitro group (dihydrogen, palladium on carbon), followed by protection of the resulting primary amine (di-*tert*-butyl dicarbonate) and oxidation of the primary alcohol (2-iodoxybenzoic acid, IBX) provided the aldehyde **34** (46%, three steps). The silyl cyanohydrin **35** was generated in 80% yield by stirring **24** with excess *tert*-butyldimethylsilyl cyanide (TBSCN) and imidazole. The benzoin addition was achieved by deprotonation of **35** with lithium di-*iso*-propyl amide (LDA), followed by addition of the aldehyde **36** (*31*). Under these conditions the α-silyloxy ketone **37** deriving from 1,2-addition, 1,3-Brook rearrangement, and cyanide elimination was obtained in 53% yield. The carbamate protecting groups were removed by treatment with hydrochloric acid (HCl) to form the diammonium salt **26**.

Silver-mediated coupling of the diamine **26** with an excess (3.9 equiv) of the β-ketothioester **25** provided the expected two-fold coupling product (LC/MS). However, all attempts to purify this product resulted in extensive decomposition deriving from cleavage of the C36–C37 bond (LC/MS analysis). To circumvent this, we developed conditions to protect this residue in situ. Thus, immediately following completion of the fragment coupling, the product mixture was transferred to a separate flask containing *tert*-butyldimethylsilyl chloride (TBSCl) and imidazole, resulting in formation of the enedisilylether **24**. The stereochemistry of the centeral alkene was determined to be (*E*), as shown, by 2D-ROESY analysis. The yield of this two-fold coupling–protection sequence was 17% (based on ^1^H NMR analysis of the unpurified product mixture using an internal standard), and **24** was isolated in 8.5% yield following reverse phase HPLC purification. By this approach, 5 mg batches of **24** were readily-prepared.

Conversion of the protected intermediate **24** to colibactin (**17**) proved to be challenging since we found that introduction of the C36–C37 α-dicarbonyl rendered the intermediates exceedingly unstable. This is consistent with an earlier model study (*31*) demonstrating rupture of the C36–C37 bond under slightly basic conditions. Ultimately, we found that treatment with concentrated hydrochloric acid in ethanol resulted in instantaneous cleavage of the carbamate protecting groups and one silyl ether; this was followed by slower and sequential cleavage of the remaining silyl ether, and aerobic oxidation to the α-dicarbonyl **38**. The α-dicarbonyl **38** was accompanied by variable amounts of the diketone hydrate (LC/MS analysis; Fig. S17 and Table S51) as observed for the bis(adenine) adduct **9** and colibactin (**17**, vide supra).

On dissolving **38** in aqueous citric acid buffer (pH = 5.0) we observed double cyclodehydration to form colibactin (**17**) and the corresponding hydrate **S33** (Fig. 7A). This mild cyclization is consistent with earlier studies establishing synthetic iminium ions resembling **38** cyclize to spirocyclopropyldihydro-2-pyrrolone genotoxins instantaneously under aqueous conditions (*22, 39*), and genetic studies supporting the off-loading of linear biosynthetic intermediates, followed by spontaneous transformation to the unsaturated imine electrophile (*23*). Synthetic colibactin (**17**) and the corresponding hydrate (**S33**) were indistinguishable from natural material by tandem MS analysis using a range of collision energies (20–50 eV, Fig. 7B and Fig. S16).

**Fig. 7.**
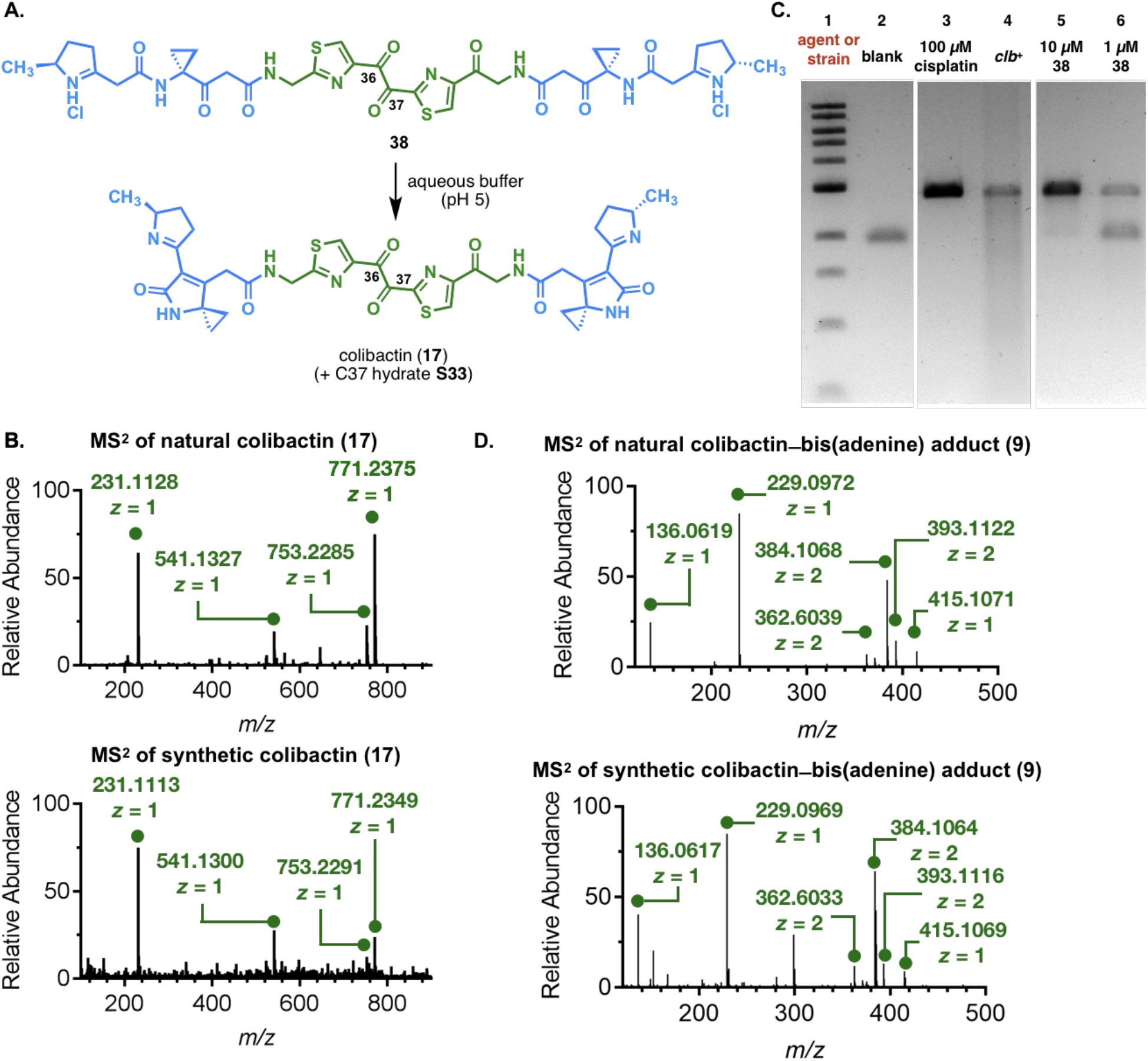
Confirmation of the predicted structure of colibactin (17). **A.** Cyclization of intermediate **38** to colibactin (**17**) and the corresponding hydrate **S33** (not shown). **B.** Tandem MS data of natural colibactin (17, top) and synthetic colibactin (**17**, bottom). Collision energy = 30 eV. For additional data see Fig. S16. C. DNA crosslinking assay employing linearized pUC19 DNA and synthetic intermediate **38.** 5% DMSO was used as vehicle. Cisplatin and *clb^+^* BW25113 *E. coli* were used as positive controls for interstrand crosslinking. DNA ladder (Lane #1); no treatment (Lane #2); cisplatin (100 μM, Lane #3); *clb^+^* BW25113 *E. coli* (Lane #4); 10 μM **38** (Lane #5); 1 μM **38** (Lane #6). Absolute concentrations of **38** are approximate. Conditions (Lane #2): linearized pUC19 DNA, 10 mM citric buffer, pH 5.0, 3 h, 37 °C. Conditions (Lane #3): linearized pUC19 DNA (15.4 μM in base pairs), cisplatin (100 μM), 10 mM citric buffer, pH 5.0, 3 h, 37 °C. Conditions (Lane #4): linearized pUC19 DNA (15.4 μM in base pairs), *clb^+^* BW25113 *E. coli*, modified M9-CA medium, 4.5 h, 37 °C. Conditions (Lanes #5 and #6): linearized pUC19 DNA (15.4 μM in base pairs), **38** (nominally 10 μM or 1 μM), 5% DMSO, 10 mM citric buffer, pH 5.0, 3 h, 37 °C. The DNA was analyzed by 0.2% denaturing agarose gel electrophoresis (90 V, 1.5 h). Additional data and controls are shown in Figs. S19, S22. **D.** (Top) MS^2^ of colibactin–adenine adduct (**9**) derived from linearized pUC19 DNA that had been exposed to *clb^+^* BW25113 *E. coli* and modified M9-CA medium, and digested. (Bottom) MS^2^ of colibactin–adenine adduct (**9**) derived from linearized pUC19 DNA that had been exposed to synthetic intermediate **38** (25 μM) at pH 5.0, and digested. Absolute concentrations of **38** are approximate.

While we could enhance mass spectral detection of natural colibactin (**17**) in the *clbS* mutant of Nissle 1917, the titers remained too low to facilitate isolation. Consequently, we turned to functional analysis of synthetic colibactin (**17**) in the DNA cross-linking assay used to detect the natural bis(adenine) adduct **9** to further confirm the structural assignment. We observed dose-dependent cross-linking of DNA (Fig. 7C) by forming colibactin (**17**) in situ from the iminium diion **38** at pH 5 in the presence of DNA. Additionally, the DNA cross-links induced by synthetic colibactin (**17**) were indistinguishable from those produced by *clb*^+^ *E. coli* (Figs. S19–S22) under basic denaturing conditions. Cross-linking was strongest at pH 5.0, and diminished as the pH was increased (Fig. S23), an observation consistent with the known instability of the α-diketone under basic conditions (*31*). An unequivocal link between synthetic and natural material was established when DNA cross-links derived from **38** were isolated, digested, and subjected to tandem MS using the same parameters employed to analyze the natural colibactin–bis(adeinine) adduct (Fig. S24). As shown in Fig. 7D (see also Figs. S112–S131 and Tables S52–S55) these tandem MS spectra were indistinguishable from the crosslinking/alkylation products derived from *clb*^+^ BW25113 *E. coli*. Collectively, the abundance of genetics data as well as these synthetic efforts confirm the structure of colibactin as **17**.

The methods used herein to elucidate the structure of colibactin (**17**) illustrate the changing landscape of natural products chemistry. As demonstrated, a combination of bioinformatics, genetics, metabolomics, and synthesis can overcome the obstacles inherent to the classical natural products paradigm of activity-guided isolation, followed by standard spectroscopic characterization. In contrast to the classical approach, by bringing a wide range of parallel experimental techniques to bear on natural product structure elucidation, it is possible to identify and characterize the activity of metastable, low-abundance metabolites. We believe that this approach will serve as a template for the identification of other non-isolable natural products, especially those formed in complex backgrounds such as the human microbiome.

## Acknowledgments

**Funding:** Financial support from the National Institutes of Health (R01GM110506 to S.B.H., 1DP2-CA186575 to J.M.C., R01CA215553 to S.B.H. and J.M.C.), the Chemistry Biology Interface Training Program (T32GM067543 to K.M.W.), the Charles H. Revson foundation (postdoctoral fellowship to A.R.H.), the NSF graduate research fellowship program (E.E.S), and Yale University is gratefully acknowledged.;

## Author contributions

M.X. characterized the natural colibactin–diadenine adduct **9** and conducted tandem MS analysis of synthetic colibactin–DNA adducts; C.S.K. characterized natural colibactin (**17**) and precolibactin 1489 (**18**) in bacterial extracts; A.R.H. contributed to the conception of the synthesis, conducted preliminary synthetic studies, and suggested protection of the fragment coupling product **24** as its enoxysilane; K.M.W. conceived the two-fold coupling approach to colibactin, developed a synthesis of the β-ketothioeseter **25**, and optimized the synthetic route; Z.W. extensively optimized the synthetic route and completed the synthesis of colibactin; M.C.F. developed a scalable synthetic route to the α-nitroketone **33**; E.E.S. generated new strains and developed the *clbS* mutant strategy to enhance detection of natural colibactin; W.W. assisted with tandem MS analysis of DNA-colibactin adducts. S.B.H. and J.M.C. conceived the study, oversaw experiments, and wrote the manuscript.

